# Connectivity by the Frontal Aslant Tract (FAT) explains local functional specialization of the superior and inferior frontal gyri in humans while choosing predictive over reactive strategies: a tractography-guided TMS study

**DOI:** 10.1101/2022.05.20.492791

**Authors:** Marco Tagliaferri, Davide Giampiccolo, Sara Parmigiani, Gabriele Amorosino, Paolo Avesani, Luigi Cattaneo

## Abstract

Predictive and reactive behaviors represent two mutually exclusive strategies for successfully completing a sensorimotor task. It is thought that predictive actions are based on the medial premotor system, in the superior frontal gyrus (SFG) and reactive stimulus-response behaviors rely on a lateral premotor system, in the inferior frontal gyrus (IFG). The frontal aslant tract (FAT), a white matter tract connecting SFG and IFG, is a possible neural substrate of the predictive/reactive interactions. We used diffusion-weighted imaging (DWI) of 17 male and female healthy human volunteers, to dissect 3 sub-bundles of fibers belonging to the left FAT (bundles 1, 2 and 3), arising ventrally from 1) the ventral precentral gyrus, 2) midway between the PCG and pars opercularis (POp) and 3) the POp and terminating dorsally in 3 different parts of the SFG, in a caudal-rostral order. We applied online transcranial magnetic stimulation (TMS) to 6 spots, corresponding to the medial and lateral terminations of bundles 1-3 during the fixed-duration set period of a delayed reaction task, that can be solved using a predictive (anticipatory) strategy or with a reactive strategy. Results showed that TMS changed the frequency of predictive/reactive strategies only when applied over 2 spots, the SFG and IFG terminations of bundle 2. Importantly, the effects of TMS were opposite when applied to the SFG or to the IFG. Our data show that the SFG and the IFG have opposite roles in producing predictive or reactive behavior and that reciprocal integration or competition is probably mediated by the FAT.

**Significance Statement:** As is well-known by athletes at starting blocks, interaction with the world can occur with a predictive strategy (anticipating a GO-signal) or a reactive strategy (waiting for the GO-signal to be manifest) and they are mutually exclusive. Here we showed, by using non-invasive brain stimulation (TMS), that two specific cortical regions in the superior frontal gyrus (SFG) and the inferior frontal gyrus (IFG) have opposite roles in facilitating a predictive or a reactive strategy. Importantly these two very distant regions but with highly interconnected functions are specifically connected by a small white matter bundle, which probably mediates the competition between predictive and reactive strategies. More generally, we show that the implementing anatomical connectivity in TMS studies strongly reduces spatial noise.

## INTRODUCTION

Whenever we must move as fast as possible in response to a sensory cue, we can adopt two different and mutually exclusive strategies. The reactive strategy consists in waiting for the GO-signal, processing bottom-up its sensory characteristics and making a movement decision as soon as the signal is detected. The predictive strategy is based on estimating the time of onset of the GO-signal and moving without waiting for the actual sensory neural information on the cue to have reached the frontal cortex. As any athlete on a starting block knows well, in the trade-off between speed and accuracy, predictive strategies are faster, but more prone to anticipation errors, while reactive strategies are slower, but more accurate.

### The reactive modality

consists in the implementation of an arbitrary sensorimotor association/rule (Passingham and Wise, 2012). The neural bases of the sensorimotor associations have been the object of several functional investigations in human brains, using complex tasks, in which a variety of prefrontal regions were identified. Imaging studies indicate arbitrary sensorimotor associations to be linked to the inferior frontal gyrus (IFG) (Rowe et al., 2000; Toni et al., 2001, 2002) and in the dorsal premotor region, around the junction between the superior frontal sulcus and the dorsal segment of the precentral sulcus with involvement of the regions of the medial wall (cingulate cortex) in the processes of outcome monitoring (Eliassen et al., 2003).

### The predictive modality

is a form of internally generated behavior. It is hypothesized that such capacity relies on a medial frontal system. Motivated, internally-generated goals are set in the medial prefrontal cortex (MPFC) and find a motor outlet in the motor cortices of the medial wall of the hemispheres, i.e. the pre-supplementary motor area (pre-SMA) and the supplementary motor area (SMA), from hereon referred to as the Supplementary Motor complex (SM-complex) (Passingham and Wise, 2012). The medial prefrontal/premotor system referred to as the “hot” prefrontal cortex (O’Reilly, 2010), the “energization system” (Stuss, 2011) or the system for “self-generated actions” (Brass and Haggard, 2008; Passingham et al., 2010; Zapparoli et al., 2017). Interestingly, the portion of cortex adjacent to the SMA and pre-SMA, on the convexity of the superior frontal gyrus seems to have functions that are assimilable to those that constitute the functional fingerprint of the SM-complex, i.e. internally generated rhythmic patterns such as finger tapping and timing-dependent actions, such as musical rhythms. The convexity of the SFG has also been found to be functionally associated to the SM-complex in the clinical manifestations of SMA lesions (Giampiccolo et al., 2021).

### The frontal aslant tract (FAT)

is a white matter tract that connects the SFG, with the ventral premotor cortex (Brodmann’ area 6 – BA6), and pars opercularis (BA 44) and pars triangularis (BA 45) of the IFG. The term “aslant” is referred to a typical oblique tract characteristic of this white matter tract that runs from the medial-superior to the inferior-lateral region (Catani et al., 2012). The FAT has been hypothesized to be involved in a series of functions related to language processes, both in the phonological and lexical domain (Kronfeld-Duenias et al., 2016; Chernoff et al., 2019; Cipolotti et al., 2020), but also in motor and cognitive functions such as verbal (Rizio and Diaz, 2016) and visuospatial (Motomura et al., 2018) working memory functions, social communication processes (Catani and Bambini, 2014), rhythm and music processing (Catani et al., n.d.; Hyde et al., 2011), and attentional processes (Garic et al., 2017). We hypothesize that such a variety of functional involvement is supported by an **underlying domain-general function that is the coordination between top-down internally generated, predictive behavior with**

### bottom-up, externally triggered actions

#### Aims of the present work

To demonstrate this hypothesis, we adopted a task that has been shown to assess simultaneously the propensity to adopt top-down predictive strategies vs. bottom-up reactive strategies, i.e., a pre-cued reaction task. This is similar to the condition in which an athlete at the starting block is informed of the upcoming GO-signal by the SET-period. In such pre-cued reaction task, two possible and mutually exclusive strategies can be implemented: anticipatory and reactive. Anticipatory behavior requires the possibility to make prior assumptions on timing of a set period (Schmidt, 1968). In one previous study from our group, (Cattaneo and Parmigiani, 2021a) a simple, built specific task showed to be appropriate at evaluating the individual propensity to act in a predictive or reactive way. This task consists in a simple delayed reaction time with a fixed and predictable set period. We observed that every subject systematically showed a biphasic distribution of response times (RTs). The first peak was too early (90 ms) to be a reactive response It was hypothesized to be due to an estimation of the SET duration, and prediction of the GO signal. The second peak (250 ms) is conversely the expression of reactive behavior, in which the subjects waits until the perception of the GO-signal and produces a response. We observed that TMS applied over a specific part of the SFG during the SET period induced a significant shift from reactive to predictive behavior. We concluded that: a) TMS is producing a “gain of function” pattern, boosting the local cortical function of SFG, i.e., predictive behavior; b) predictive and reactive strategies are processed in a parallel way and therefore are available for selection until the very final stages of motor preparation and c) TMS has been able to temporally impair the balance between the two behaviors. According to our model that postulates the SFG and IFG having opposite roles in promoting predictive and reactive behavior respectively. Therefore, in the present experiment, we hypothesized that applying TMS to the SFG or the IFG should produce a double-dissociation pattern, with IFG stimulation leading to a preference for reactive behavior and SFG stimulation leading, as already demonstrated to a shift towards predictive behavior. Moreover, we hypothesized that anatomical connectivity by FAT fibers should be a spatial constraint in indicating the specific regions of the SFG and IFG where the double dissociation pattern was to be expected.

## METHODS AND MATERIALS

### Subjects and MR session

17 subjects participated in the study (8 females and 9 males). The experiment was approved by the local Ethical Committee of the University of Trento (protocol 2020_035) and all participants signed informed consent papers to join the study. Participants were screened for TMS contraindications prior to the experiment (Rossi et al., 2009, 2011). All participants were required to join two separate sessions: the first one was an MRI-DTI session and the second one a neuronavigated TMS stimulation session during the performing of a task. The anatomical images were acquired by a 3T MAGNETOM Prisma (Siemens Healthcare, Erlangen, Germany) with a 64-channel head-neck RF receive coil was used to acquire 3D T1-weighted (T1w, multi-echo-MPRAGE, 1mm-isotropic), and diffusion-weighted (dMRI) data (2mm-isotropic, TE/TR=76/4200ms, shells: b={0,700,1000,2850} s/mm, 32/64/64 directions as described in Hubner et al. (2022). DW imges processing. T1-w images of all the subjects underwent a standard preprocessing. First of all, the raw DICOM T1-w images were converted to NifTI format using dcm2niix software (Li et al., 2016). Then, an AC-PC (Anterior-Posterior Commissures) alignment was performed by means of rigid registration to the MNI152 T1-w template (Avants et al., 2008). The brain mask was estimated using a pre-trained 3D U-Net (Ronneberger et al., 2015; Çiçek et al., 2016). The brain mask was used to perform the Bias-Field Correction restricted on the brain voxels, using N4-Bias Field Correction tool (Tustison et al., 2010). Furthermore, the T1-w images were segmented into 6 brain tissue by means of a pre-trained 3D-Unet (Çiçek et al., 2016; Amorosino et al., 2020). To support the alignment of structural and diffusion images we computed a synthetic T2-w image using AFNI toolkit (Cox, 1996).

#### DWI data preprocessing

The preprocessing of the Diffusion-Weighted Imaging (DWI) data was carried out using TORTOISE toolkit (Pierpaoli et al., n.d.). With DIFFPREP tool (Okan et al., n.d.) we computed the correction for Gibbs ringing, thermal noise, eddy current, and motion distortion. Furthermore, we used DRBUDDI tool (Irfanoglu et al., 2015) to perform the susceptibility induced echo-planar imaging (EPI) distortion correction through diffeomorphic registration (Avants et al., 2008) of the DWIs volumes. DRBUDDI was operated combining both DWI reversed-phase encoding directions (i.e. AP and PA acquisitions) and also the information from an undistorted structural MRI.The DWIs data were also corrected for bias field inhomogeneities (Tustison et al., 2010).

The reconstruction of the streamline tractography was performed using MRtrix3 software (Tournier et al., 2012). We performed the multi-shell, multi-tissue Constrained Spherical Deconvolution (CSD) (Jeurissen et al., 2014) (lmax=6; threshold=0.5) to obtain the WM fiber Orientation Distribution Function (fODF) from the estimation of the response function using the Dhollander method (Dhollander et al., 2016; Raffelt and Connelly, 2019). Then, we computed the deterministic tractography based on CSD (cutoff 0.001, maximum angle of 75 degrees, step of 0.5 mm) constraining the length of the streamlines between 20mm and 250mm. We initialized the tractography using random seeding of 10^7 seeds on a WM mask, estimated by thresholding the DTI Fractional Anisotropy scalar map with a heuristic value of 0.15. We stopped the tracking by selecting 2*10^6 streamlines.

#### Individual FAT dissection and neuronavigation

Once the full masked tractogram was obtained, it was processed in TrackVis to dissect the left FAT portion of interest. Note that while there is consensus on the caudal limit of the FAT from the ventral precentral gyrus, the anatomical definition of the FAT lacks a clear rostral border (Varriano et al., 2018). Therefore, a rostral limit was set in the present work to a region of interest i.e., the group of fibers connecting the SFG with the ipsilateral ventral precentral gyrus and IFG - pars opercularis (***Figure 1A***). The seed ROIs in the SFG and IFG were set manually, following individual anatomies. False positive tracks were excluded by applying a band-pass length filter (50-90 mm), a x-plane filter, excluding fibers that crossed the midline, a y-plane filter excluding fibers caudal to the central sulcus and finally by means of manual removal upon visual inspection using the Tractome toolbox (Porro-Muñoz et al. 2015). The left FATs were then divided into 3 equal sub-bundles, hereafter named posterior, middle and anterior (***Figure 1B***). The 6 dorsal and lateral cortical origins of the 3 sub-bundles were then used as targets for neuronavigated TMS by means of a neuronavigational system (SofTaxic software v.3.4, EMS, Italy), labelled as points from P01 to P06 with odd points in the dorsal origins and even points in the medial origins. For the purpose of further analysis, the 6 points were treated as three pairs of homologous points, one dorsal and one ventral and each connected by a single sub-bundle (***Figure 1C***). Individual T1-weighd MRs were used for stereotaxic frameless neuronavigation, by means of an optical infrared camera and reflective markers placed on the participant’s head and TMS coil.

**Figure 1.**
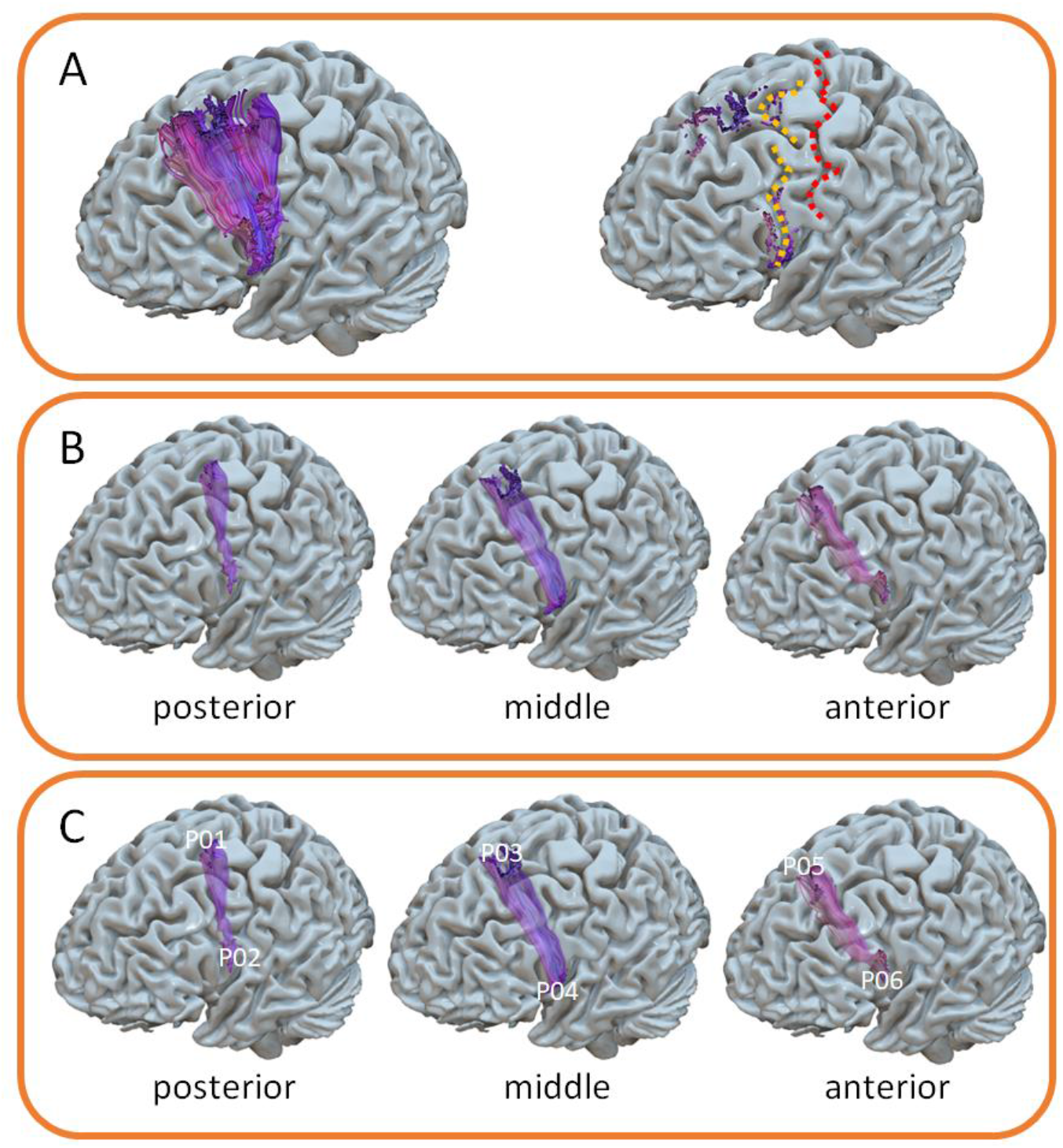
**A**. Left FAT in a representative paritcipant. The left image shows the central sulcus (in red) and the precentral sulcus (in yellow). **B**. division of the left FAT in 3 sub-bundles. **C**. identificationo of the 3 pairs of homologue cortical targets for TMS.

#### TMS

We used a MagPro 100 stimulator, connected to a figure-of-eight TMS coil with 65 mm diameter of each winding (model MagPro MCF-B65). The individual resting motor threshold (rMTh) was assessed in the left hemisphere by stimulating the optimal spot for the visible contraction at rest of the right *Opponens Pollicis* muscle (OP). RMTh was assessed as the minimum stimulation intensity that elicits a visible contraction in intrinsic hand muscles with a 50% probability over a series of 10 stimuli. The stimulation intensity for the experiment was then set to 120% of the individual rMTh. Sham stimulation was achieved by tilting the coil by 90° to the scalp surface.

#### Experimental procedure and task

Once the neuronavigational tools were calibrated, 3D-model of the subject’s brain with ROIs applied was prepared and stimulation intensity was obtained, subjects were required to place their head on a fixed chinrest for the entire length of the experiment at 60cm from a 27’’ standard monitor with a refresh rate of 60Hz, and a keyboard placed in front of them. Stimulus presentation and synchronization with TMS triggering was performed using MATLAB (Psychtoolbox) scripts and delivered by MATLAB v.2018b. The experimental task consisted in a pre-cued GO-task, i.e., a sequence of ‘rest’, ‘set- period’ and ‘GO+response’. A schematization of a single trial is shown in ***Figure 2***. Participants were warned of the upcoming GO-signal by the set period, which had a fixed duration (800 ms). The task was similar to that used in (Cattaneo and Parmigiani, 2021b). The color of a circle in the center of a white screen informed the Participant of the phase of the trial: grey for rest, yellow for the set-period, and green for the GO-signal. The instruction was to press the spacebar with the right hand as fast as possible. Experimental instructions included the information that the duration of the set-period was fixed. Feedback was given when the button was pressed before the GO-signal (false start) in the form of the words “too early” on the screen and when the button was pressed too late “too late”. The whole script for stimulus presentation and TMS triggering is available as supplementary material at: https://osf.io/x76cm/. The grey circle duration (rest phase) varied randomly according to a square-wave function, between 1000 ms and 3000 ms. The yellow circle (set-period) persisted for a constant time duration of 800ms and the GO-period of 600ms for the green one. The coil of the TMS was placed tangentially on the relative left FAT endpoint selected on the T1 image, with a local error less than 1.7 mm and TMS was delivered at a random time at 400-800ms from the onset of the set-period. TMS was delivered to the left hemisphere, contralateral to the responding (right) hand. Participants wore earphones for the entire duration of the experiment, listening to white noise interspersed with sounds of TMS stimulation, generated by the MATLAB script TMSNOISE (Russo et al., 2022).

**Figure 2:**
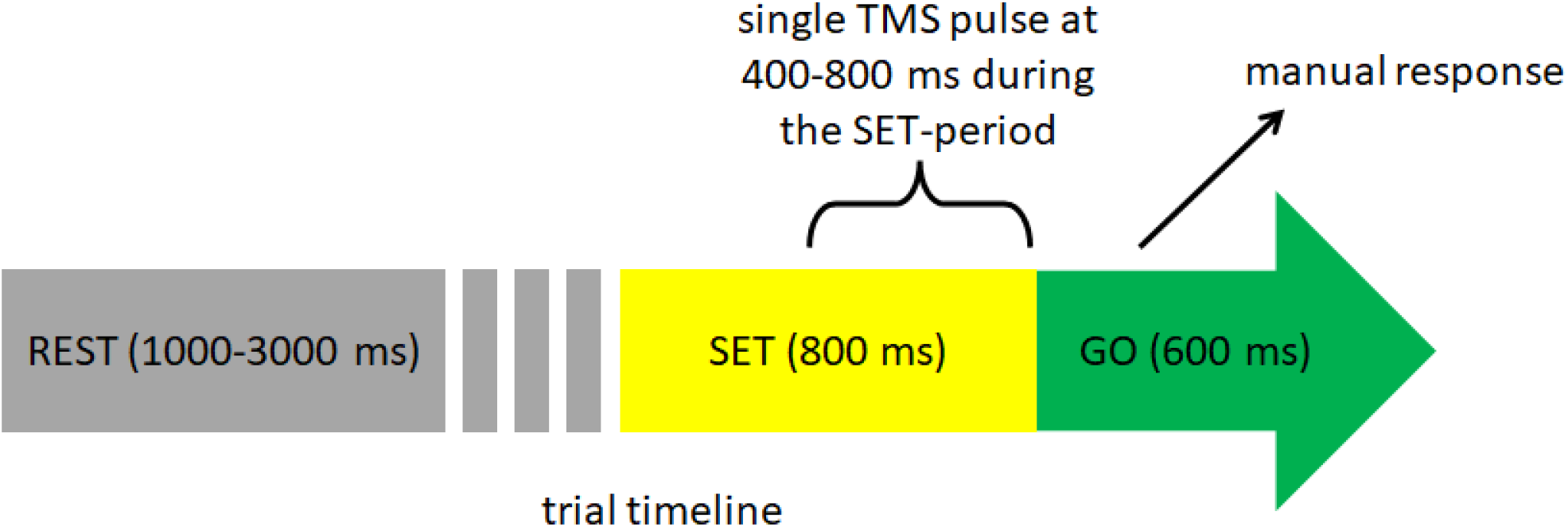
schematization of a single trial. TMS was delivered at a random time /square wave function) comprised between 400 and 800 ms form the beginning of the set-period.

#### Experimental design

The experiment had a within-subjects design, in which the dependent variable was a measure of the frequency of reactive behaviour (the Reactive Index, see below) and the independent variables were the stimulation sites. We employed a blocked design in which a single stimulation site was targeted in a single block of 40 trials. The order of the 6 blocks (and therefore of stimulation sites) was randomized between participants. In addition, we ran 3 extra blocks with sham stimulation, which occurred always at the beginning, in the middle and at the end of the sequence of blocks. This accounted for a total of 40 trials (one block) for each active TMS site, and 120 trials (3 blocks) for sham TMS. Before the actual experiment the participants familiarized with the task in a series of at least one practice block, until they reported confidence with the task.

#### Data processing

The response time (RT) was calculated as difference between the time of response and the GO-signal and therefore could have both negative and positive values. RT was the main experimental output and index of performance, because it allowed to classify trials as product of a reactive strategy or of a predictive strategy. Preliminary data cleaning was done with the purpose of excluding trials in which the response had been given too early to be influenced by TMS (exclusion of trials with response occurring earlier than 50 ms from TMS) and to exclude trials in which the participant lingered excessively, thus not fulfilling the criterion for responding as fast as possible (trials with RTs > 600 ms). At this point, trials were classified as A) false starts (i.e., trials with response before the GO-signal, i.e., negative RTs); B) anticipatory trials (i.e., trials with RT between 0 and 170 ms) and C) reactive trials (trials with RTs > 170 ms). The value of 170 ms was chosen empirically by inspecting the average trough in the two peaks of the RT distribution, as also performed in (Cattaneo and Parmigiani, 2021b) and confirmed by the present data, as shown here in ***Figure 3***. We then took into consideration the pool of trials within each single condition (40 trials per each active TMS site) and 120 trials for the sham, and built a ratio called “reactive index” (RI), by dividing the number of reactive trials by the total number of trials.

**Figure 3:**
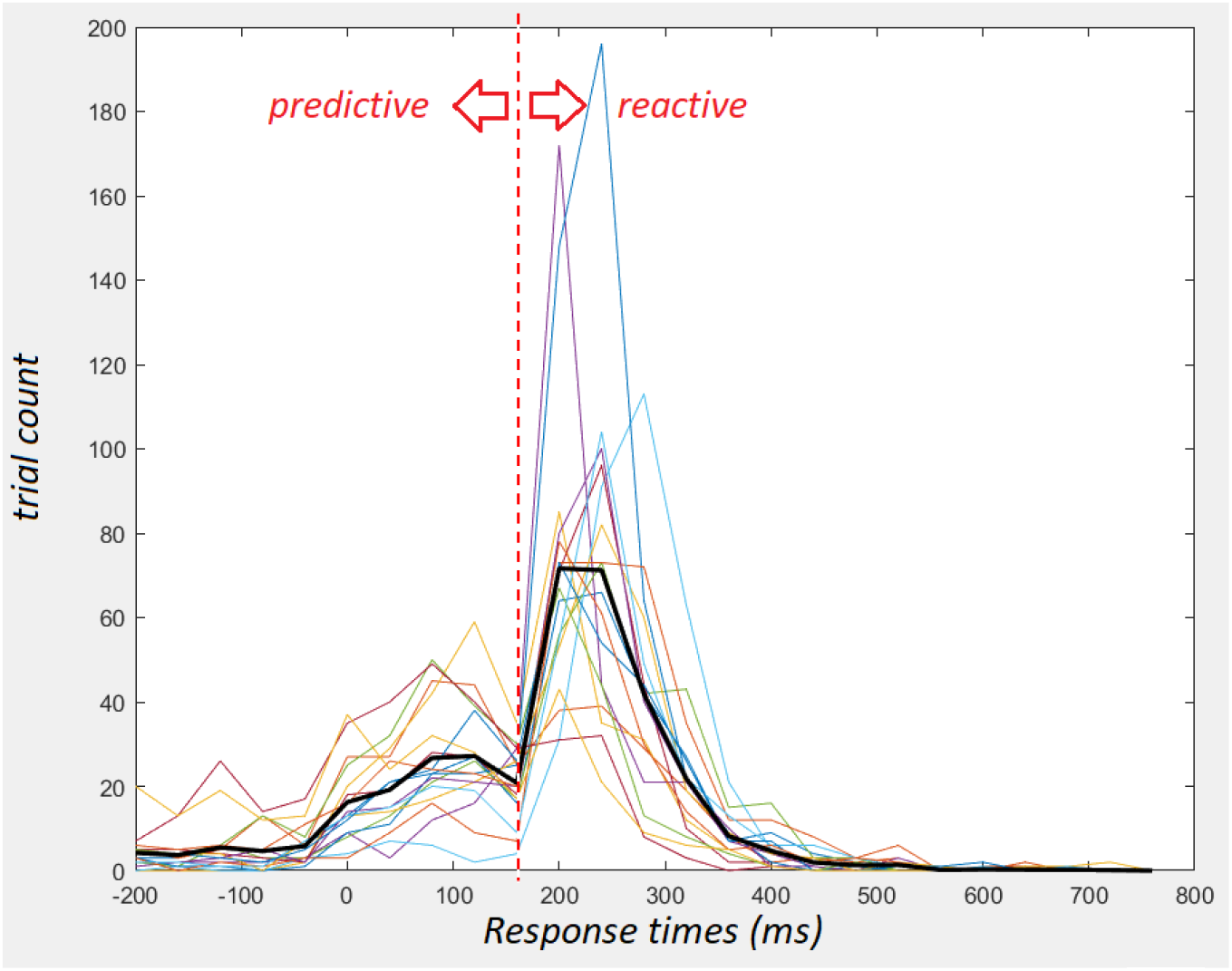
distribution of individual response times of all 17 participants over 40 ms bins (thin colored lines). The average value of the population is shown by the thick black line.

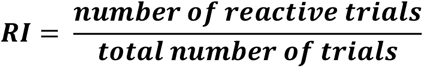

The RI is distributed between 0 (100% of predictive behavior) and 1 (100% of reactive trials). In this way the data from all participants were reduced to 7 data points per subject: the RI from the 6 active TMS sites and the RI from the sham condition. Finally, further data reduction was obtained by subtracting the RI from Sham TMS from the 6 RI values of the active TMS spot. The differential RI obtained was a number distributed between -1 and +1 and indicated, if negative, that effective TMS had increased the number of predictive trials compared to Sham and, when positive, that TMS had increased the number of reactive trials compared to Sham. The differential RIs were the actual variable that was used in the statistical analysis

#### Statistical analysis

The aim of statistical analysis was to assess whether TMS over a specific site significantly biased the propensity to act with a reactive or a predictive strategy compared to other TMS sites. The 6 sites were clustered according to 2 orthogonal dimensions: the medial-lateral (M-L) dimension and the caudal-rostral (C-R) dimension. We therefore performed an ANOVA on the differential R-I as dependent variable and with 2 within-subjects factors: M-L (2 levels= medial and lateral) and C-R (3 levels = posterior middle and anterior). The second aim was to understand whether, in absolute terms, each single active TMS site was associated with an absolute increase or decrease in the propensity to act in predictive or reactive way. This was demonstrated by testing the means of the differential-RI of each of the 6 spots compared to the null hypothesis of mean=0 (i.e., no difference compared to sham). In this case, given the univariate approach, we adjusted the significance value to a more conservative threshold d of p = 0.01.

#### Data availability

The whole dataset of anatomical data and behavioral data are accessible together with the software used for data analysis at the following repository: https://osf.io/x76cm/

## RESULTS

None of the participants reported any significant immediate nor delayed undesired effect of TMS. The distribution of RTs confirmed the main hypothesis behind the use of the task, i.e. the bimodal distribution of the data, with a clear trough between the two peaks that was similar across participants and centered around 170 ms. The early peak is made of predictive responses, the late peak is made of reactive responses (***Figure 3***).

The ANOVA showed marginal effects of the caudal-rostral dimension (main effect of C-R: F(2, 32)=3.23, p=0.053) and of the medial lateral dimension (main effect of the M-L dimension: F(1, 16)=3.97, p=0.063). The most important result was a interaction between the M-L and the C-R dimensions (F(2, 32)=7.85, p=0.001) associated with indicators of a large and robust effect size (eta-squared = 0.33). The interaction was further explored by breaking it down into 3 paired t-tests comparing the means of the 3 pairs of homologue spots in the SFG and IFG, i.e., P01-P02, P03-P04 and P05-P06 (see Figure 1 for the homologies between spots). The results indicated no significant difference neither for the P01-P02 pair (t(16)=0.53, p=0.61) nor for the P05-P06 pair (t(16)=-0.65, p=0.36). We observed however a significant difference between the P03 and the P04 homologues (t(16)=-4.97, p=0.0001) The data are shown in ***Figure 4***.

**Figure 4.**
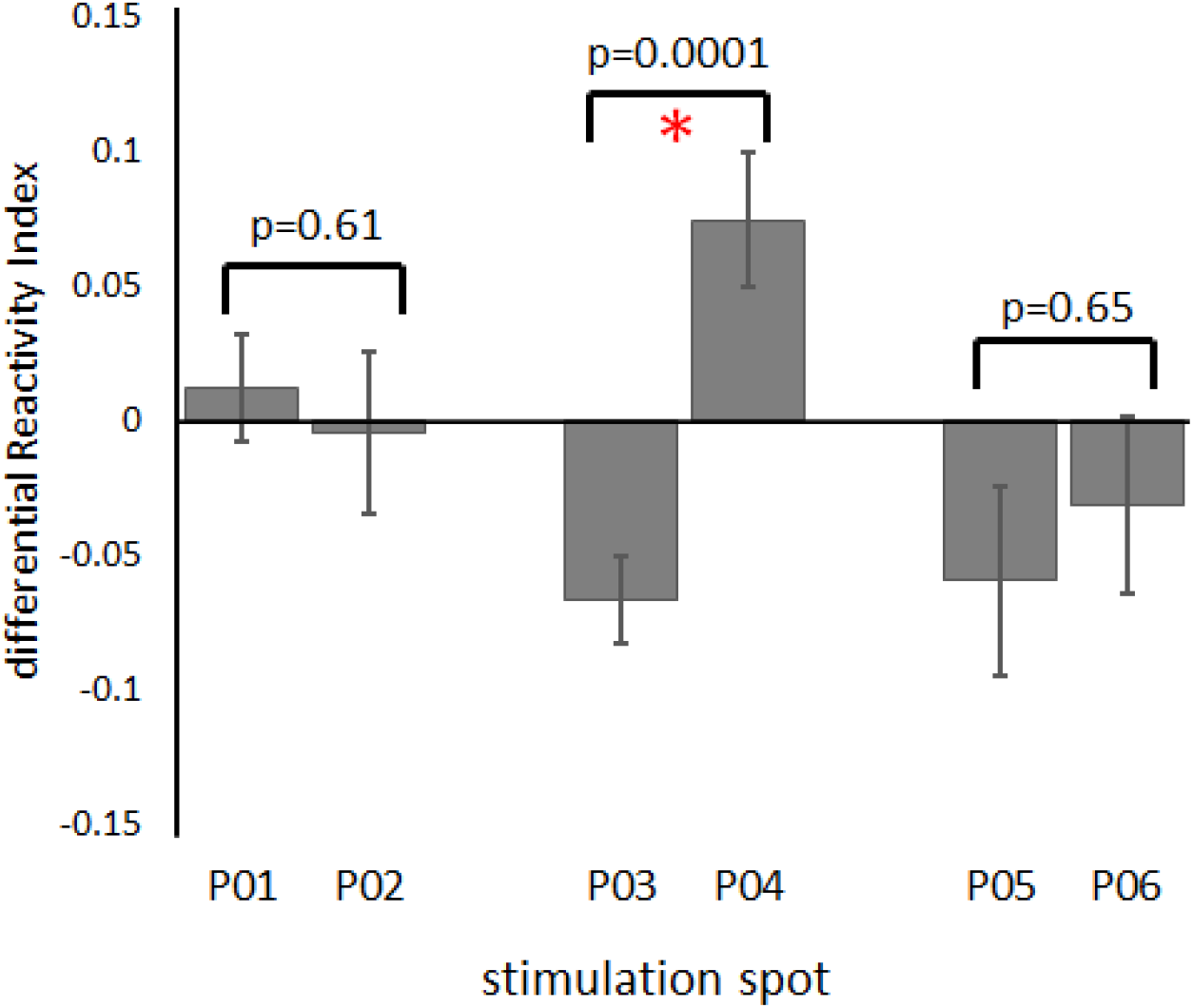
Mean values of the differential reactivity index and results of the t-statistics comparing the mean values between homologue spots along the M-L dimension (note that negative values indicate greater frequency of predictive behavior compared to Sham stimulation and positive values indicate greater frequency of reactive behavior compared to Sham).

The univariate analysis testing the difference of differential RIs against the null hypothesis of mean=0 (indicating no difference from the Sham – baseline – condition) showed that the P03 and P04 spots were significantly different from the null hypothesis. In particular, (i)**TMS over P03 induced significantly more predictive behavior** compared to Sham. (ii) **TMS over P04 induced significantly more reactive behavior** compared to Sham. The data and the statistics are presented in ***Table 1*** and ***Figure 5***

**Table 1:** mean values of the differential RI for each stimulation spot (note that negative values indicate greater frequency of predictive behavior compared to Sham stimulation and positive values indicate greater frequency of reactive behavior compared to Sham). The t-statistics for single sample against the null hypothesis of mean=0 are shown. Significant conditions are highlighted in bold.

| TMS spot | Mean | Std.Dv. | N | Std.Err. | t-value | df | p |
| --- | --- | --- | --- | --- | --- | --- | --- |
| P01 | 0.012 | 0.081 | 17 | 0.019 | 0.603 | 16 | 0.55 |
| P02 | -0.005 | 0.124 | 17 | 0.030 | -0.154 | 16 | 0.88 |
| P03 | -0.067 | 0.067 | 17 | 0.016 | -4.085 | 16 | 0.0009 |
| **P04** | **0.074** | **0.101** | **17** | **0.024** | **3.006** | **16** | **0.008** |
| P05 | -0.059 | 0.145 | 17 | 0.035 | -1.684 | 16 | 0.11 |
| P06 | -0.031 | 0.136 | 17 | 0.033 | -0.949 | 16 | 0.36 |

**Figure 5.**
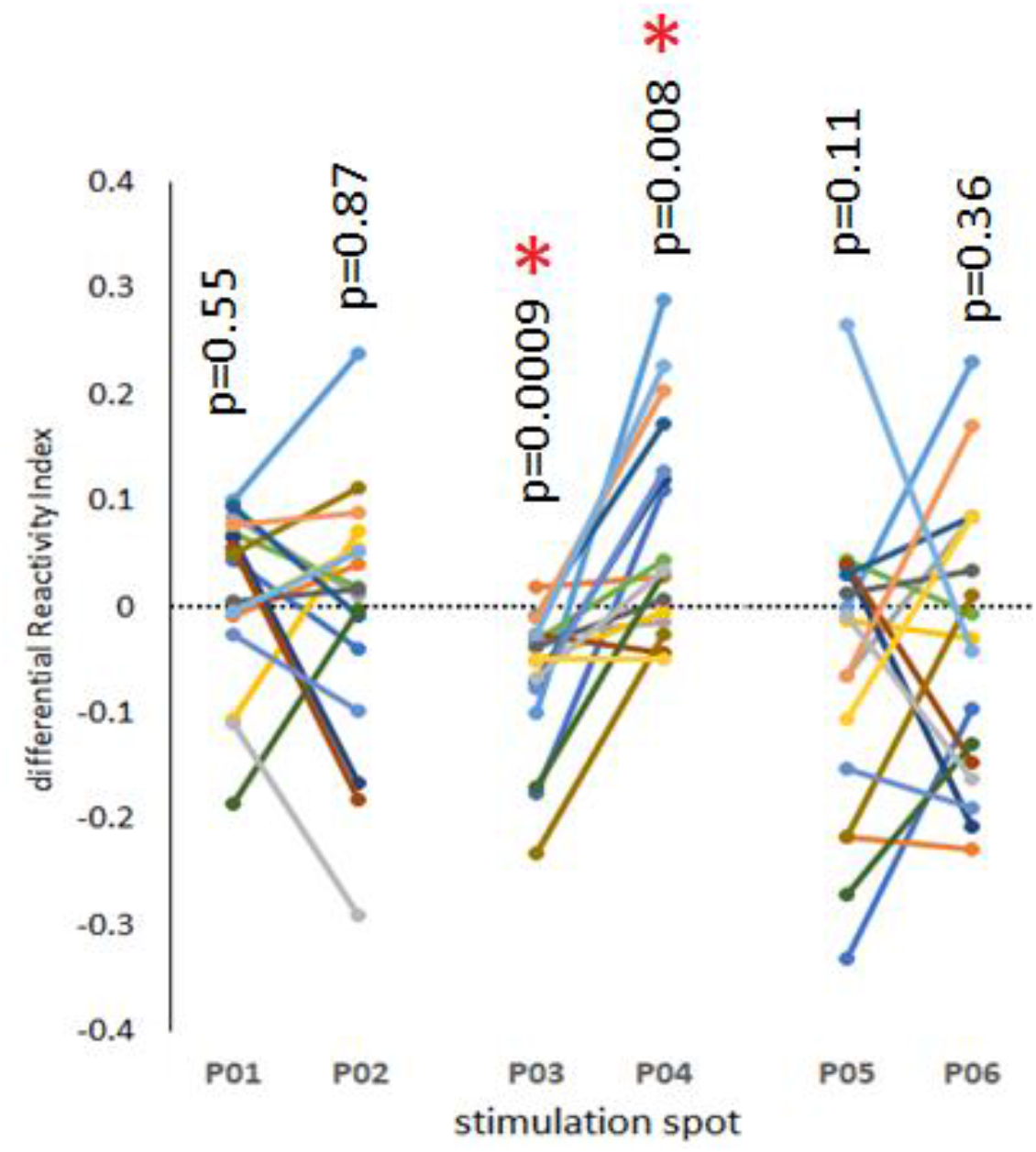
Individual values (n=17) of the differential Reactivity Index in the 6 TMS spots. The t-statistics for single sample against the null hypothesis of mean=0 are shown.

**Figure 6.**
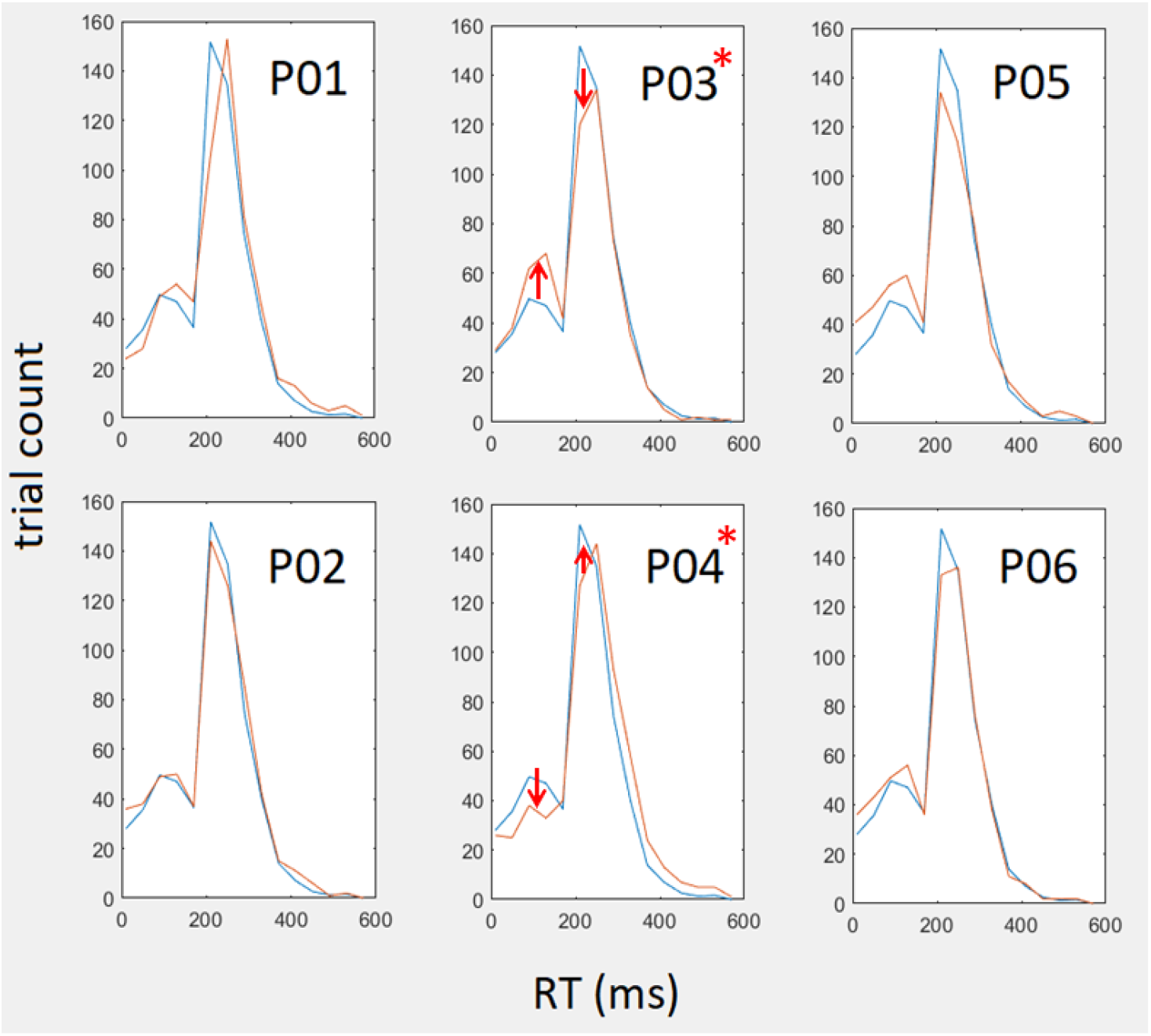
The representation of the mean distribution of response times (RTs) in the sham condition (blue line – repeated in every panel) and in the 6 active TMS spots (red lines). Red arrows and asterisks indicate the changes in the distribution of the predictive/reactive ratio in the spots where this was significant. Note that the graph shows the average values from the population and therefore is for illustrative purposes. Quantification of effects and statistical significance are to be extrapolated from Figures 4 and 5.

## DISCUSSION

### The SFG promotes internal predictive behavior and the IFG promotes reactive behavior

In the present experiment we have investigated the reciprocal role of the SFG and IFG in selecting a predictive or a reactive action strategy. Our specific hypothesis was that the SFG and IFG have reciprocally opposite roles, with the SFG promoting predictive behavior and the IFG promoting reactive behavior and that their interaction is mediated by the FAT. We employed a validated task that simulates a “starting block” scenario. Participants had to respond to a pre-cued GO-signal, preceded by a SET-period of fixed and predictable duration while TMS was delivered during the SET-period. This task can be solved in one of two mutually-exclusive strategies: the predictive and the reactive one. More in detail, stimulation of a specific portion of the SFG induced a bias towards predictive behavior, thus replicating the data from Cattaneo and Parmigiani 2021 (Cattaneo and Parmigiani, 2021a). The effective spot was the one referred to as P03 (see Figure 1 for the location of TMS spots). This result is fully consistent with the role played by the SFG in timing of action. This has been investigated mainly on the medial region of the SFG, that is, in the SM-complex, where several neural signals that are causally linked to time processing and to timing of action, at the basis of movement based on internally cued anticipatory strategy (Mauk and Buonomano, 2004; Chen et al., 2006; Mita et al., 2009; Casini and Vidal, 2011). Stimulation of the IFG, specifically of the P04 spot, produced behavioral effects opposite to those over P03, i.e., an increase the propensity to perform reactive responses. This result is coherent with current knowledge on IFG function. The IFG is known to code for sensorimotor associations in rule-dependent behavior (Toni et al., 2001). Most of the evidence in favor of such hypothesis come from studies in non-human primates, where in the sectors of F4 and F5 sensorimotor neurons have been described supporting 1:1 sensorimotor associations (Rizzolatti et al., 1998; Bonini et al., 2014), in the grasping and reaching domains (Gentilucci et al., 1983; Fogassi et al., 1996).

### Polarity of TMS effects

We operate under the assumption that TMS here had a facilitatory effect on behavior. Therefore, the effect of TMS on the SFG promoting predictive behavior indicates that the functional role of SFG is that of producing predictive behavior, and this was enhanced by single-pulse TMS. Vice-versa, the effects of TMS on the IFG, consisting in an increase in reactive behavior indicate that the role of the SFG is that of promoting reactive behavior, which was enhanced by single-pulse TMS. This is coherent with most literature on TMS-behavior relations postulating that the effects of TMS are state-dependent and cannot be predicted as loss-of-function or gain-of-function, without information on timing and intensity of TMS and most importantly, without information on the excitability state of the stimulated cortex. During these years the TMS has been described as a tool that allows us to induce a “virtual lesion” on a stimulated area, because of similarities in functioning that we can see between a lesioned area and an area stimulated by the TMS. This description over the years has been discussed and integrated with other definitions. It is preferable to describe the effects of TMS as dependent of the subject’s brain condition during the stimulation (Silvanto and Muggleton, 2008). Generally we define the effects of TMS in four ways: virtual lesions (TMS suppresses signals), noise generation (TMS intensity-dependent), activity-related enhancing (TMS generates random activity) and mediating the activity of neurons in the stimulated area (Perini et al., 2012). Some authors also described the TMS effect as “activity-dependent” or dependent on the “initial neural activation state”, meaning that manipulating neural activity before TMS delivering It is possible to select specific neural populations to stimulate within the area we are interested in (Silvanto et al., 2008; DS et al., 2011; Perini et al., 2012)

### Coexistence of different motor programs and parallel processing of information

A recent study by our group showed that TMS during the SET period can induce biases towards one of the two strategies. (Cattaneo and Parmigiani, 2021a). This indicates that up to the reaction time, during the SET-period, both strategies are still available and present in parallel channels in the participant’s motor system. If the commitment to one of the strategies was determined earlier on during neural processing, we would not be able to induce a strategy switch with TMS during the SET period. The present data provide evidence in favor of parallel processing of the two possible strategies. The concept of parallel channels in the action system that mediate bottom-up (in this case the reactive strategy) and top-down (the predictive strategy in our protocol), that compete for motor output, is well established in several models of the action system (Kornblum et al., 1990; Ridderinkhof et al., 2004; Cisek and Kalaska, 2010; McBride et al., 2012). Several lines of empirical evidence in human neurophysiology (Michelet et al., 2010; Barchiesi and Cattaneo, 2013a; Ubaldi et al., 2015) and human behavior (Van Zoest and Donk, 2006; Barchiesi and Cattaneo, 2015) confirm that top-down control and bottom-up sensorimotor processes seem to coexist up to the very distal phases of action production.

### Anatomical connectivity explains the effects of TMS over distant but interconnected regions

According to our experimental hypothesis, the FAT provides the anatomical substrate for interaction between the SFG and the IFG. In the context of this model, we hypothesized that specific sectors of the SFG and the IFG influencing the propensity to act in a predictive or reactive way should be directly connected by FAT fibers. Indeed, we demonstrated that stimulation of two limited portions of cortex, connected by a bundle of fibers, produced related behavioral effects. The effects of stimulation of the SFG were limited to the portion of cortex that was directly connected to P03, i.e., the P04 spot. We show here that coupling different sectors of the SFG and the IFG in terms of homolog regions connected by sub-bundles of the FAT indeed explained the totality of the variance of the effects of TMS over the two gyri. What’s more relevant, is that the effects of TMS on the two FAT terminations were opposite. We hypothesize that these two regions act with a mechanism of reciprocal (mutual) inhibition. On a microscale, mutual inhibition is a widely used neural mechanism for selection between competing and mutually exclusive actions, as observed in non-human species (Machens et al., 2005; Koyama and Pujala, 2018). Based on this concept, we can find a connectivity pattern known as mutual inhibition of lateral inhibition. This phenomenon is a complex series of processes that can reinforce certain neuronal effects providing computations useful for simple and complex behavioral patterns. By these processes It is not possible to elicit two possible behavioral patterns in response to a certain situation because one excludes the other. Similarly in our study we observed not only that we have the same kind of TMS-induced interferences between homologous regions of SFG and IFG, but also that there is a certain complementarity between these regions, depending on the spot stimulated we have recollected an increase of one strategy (predictive) or the opposite strategy (reactive) between the two opposite sides of the FAT endpoints. The resulting neural decisions follow a “winner takes all” pattern, depending on the balance between IFG and SFG outputs. Summing up, we identified here a connectomic pattern in which two distant cortical regions connected with high spatial specificity by a direct white matter bundle seem to form a functional unit with the role of reciprocal competition.

### Consistence of the present data with current knowledge on the FAT

The exact function of the FAT is currently unknown. Several hypotheses have been proposed and one authoritative model postulates that the FAT connectivity plays a domain general role, in motor planning and timing of more or less complex motor sequences, in monitoring and in the interactions between executive control and sensorimotor patterns (Dick et al., 2019). This said, there is strong evidence that such domain-general role is “borrowed” specific functions in the two hemispheres. Indeed, several lines of evidence point at the concept that FAT functions may be lateralized (La Corte et al., 2021). A right-lateralized SFG-IFG circuit is specialized for inhibitory control. Studies showed that lesions over the right pars triangularis and right pars opercularis slowed down the SSRT results during STOP and GO tasks, or its involvement has been found during interference suppression, suggesting a particular role in inhibitory control or suppressing processes during task performance (Bunge et al., 2002; Aron et al., 2003). Similarly, other authors have found a predominant role of the right pre-SMA in action inhibition (Sumner et al., 2007). On the left hemisphere, the FAT has been correlated to motor-speech responses (posterior FAT) and lexical-semantic processes (anterior FAT) (Corrivetti et al., 2019), as well as lexical recalling deficits induced by lesions over left FAT, and relative endpoints on pre-SMA and SMA, based on verb generation task on awake surgery while an intraoperative electrical stimulation was applied (Sierpowska et al., 2015). Lesions on the FAT were found to be responsible of verbal fluency deficits, and a dissociation between verbal fluency and semantic functions (Catani et al., n.d.), and similarly patients with FAT disconnection experienced impairments in phonemic fluency tests (Cipolotti et al., 2020). Developmental alterations of the FAT have been associated with congenital stuttering (Kronfeld-Duenias et al., 2016). The FAT has been found to be involved in working memory functions (Varriano et al., 2018) by two-back working memory tasks, and in social communication functions (Catani and Bambini, 2014). information about homologous regions between areas connected by the FAT has not been fully covered by studies. There are studies that describe the FAT parcellation, depending on its extension respect to the SFG, for instance some authors observed that the FAT needs an extended definition since It can end further anteriorly labelling it as “ExFAT” (Rojkova et al., 2015; Pascual-Diaz et al., 2020), while other authors restricted the extension (Ferpozzi et al., 2018) originally proposed by Catani et al (Catani et al., 2012).

### Conclusions

We identified a specific fronto-frontal circuit representing a functional unit involved in mediating interactions between predictive and reactive behavior finalized towards the same goal. In response to our original scientific question and aim of the study, we shown evidence in support of the FAT as a system mediating reciprocal inhibition between two regions of the premotor cortex that code for predictive (SFG) vs reactive (IFG) strategies. This connection system supports strategy selection in a “winner takes all manner”. In addition, we show here for the first time that individual information on anatomical connectivity (tractography) coupled with TMS can significantly increase the signal to noise ratio in spatial mapping of the cerebral cortex and provides a whole new way to interpret the functional mapping of the brain by non-invasive stimulation techniques.

## Acknowledgments

We wish to thank Enrica Pierotti for the valuable help in data collection and analysis. Current affiliations for Dr Giampiccolo are: Department of Clinical and Experimental Epilepsy, UCL Queen Square Institute of Neurology, University College London, London, UK. Victor Horsley Department of Neurosurgery, National Hospital for Neurology and Neurosurgery, Queen Square, London, UK Institute of Neurosciences, Cleveland Clinic London, London, UK

